# In-Situ High-Resolution Cryo-EM Reconstructions from CEMOVIS

**DOI:** 10.1101/2025.03.29.646093

**Authors:** Johannes Elferich, Marek Kaminek, Lingli Kong, Adolfo Odriozola, Wanda Kukulski, Benoît Zuber, Nikolaus Grigorieff

## Abstract

Cryo-electron microscopy can be used to image cells and tissue at high resolution. To ensure electron transparency, sample thickness must not exceed 500 nm. Focused-ion-beam (FIB) milling has become the standard method to prepare thin samples (lamellae), however, the material removed by the milling process is lost, the imageable area is usually limited to a few square microns, and the surface layers sustain damage from the ion beam. We have examined <u>c</u>ryo-<u>e</u>lectron <u>m</u>icroscopy <u>o</u>f <u>v</u>itreous <u>s</u>ections (CEMOVIS), a preparation technique based on cutting thin sections with a knife, as an alternative to FIB-milling. CEMOVIS sections also sustain damage, including compression, shearing and cracks. However, samples can be sectioned in series, producing many orders of magnitude more imageable area compared to lamellae making CEMOVIS an alternative to FIB-milling with distinct advantages. Using 2-dimensional template matching on images of CEMOVIS sections of *Saccharomyces cerevisiae* cells, we reconstructed the 60S ribosomal subunit at near-atomic resolution, demonstrating that, in many regions of the sections, the molecular structure of these subunits is largely intact, comparable to FIB-milled lamellae.

## 1. Introduction

An important goal in cell biology is to understand how cellular function arises from the complex interactions of the molecules and their assemblies inside a cell. Using electron tomography, 3-dimensional (3D) reconstructions of the cellular environment can be obtained to visualize the spatial arrangement of molecular machines, membranes and organelles (Robinson *et al*., 2007; Mahamid *et al*., 2016). To be electron-transparent, samples have to be thin, ideally 100 – 200 nm thick. A standard technique to generate thin samples from cells and tissue is sectioning with a knife. Traditionally, sections are cut from chemically fixed and resin-embedded samples (Studer & Gnaegi, 2000; Sader *et al*., 2007). However, chemical fixation, dehydration, and embedding techniques do not preserve the structure of these samples at the molecular level (Dubochet & Sartori Blanc, 2001; Sader *et al*., 2007), leading to the development of <u>c</u>ryo-<u>e</u>lectron <u>m</u>icroscopy <u>o</u>f <u>v</u>itreous <u>s</u>ections (CEMOVIS, (Dubochet *et al*., 1983; Al-Amoudi *et al*., 2004)). To prepare samples for CEMOVIS, cells or tissues are vitrified by high-pressure freezing (Studer *et al*., 2001), a technique that preserves the sample by preventing the formation of ice crystals. Thin sections are cut from the frozen sample in a cryo-ultramicrotome, producing a ribbon of sections that can be transferred onto a grid for imaging by cryo-electron microcopy (cryo-EM) or cryo-electron tomography (cryo-ET). The integrity of the sections depends critically on the details of the cutting process, and much development has been invested in optimizing this step (Al-Amoudi *et al*., 2003; Ladinsky *et al*., 2006; Studer *et al*., 2014). Despite these efforts, cryo-EM images of sections usually show evidence of compression in the direction of the movement of the knife, as well as ridges and crevasses that may originate from the bending of the cut section as it is lifted off the bulk sample by the knife (Han *et al*., 2008; Al-Amoudi *et al*., 2005; Hsieh *et al*., 2006). Furthermore, there is shearing of the sample that leads to visible discontinuities in membranes and filaments (see below), suggesting that there is also damage on the molecular scale.

To avoid the type of sample damage seen with CEMOVIS, an alternative approach for the preparation of thin samples from cells and tissue was developed, based on the removal of sample material by a focused ion beam (FIB-milling, (Marko *et al*., 2006)). FIB-milling is now being used routinely, with dedicated instrumentation integrating light microscopy to locate areas of interest that are labeled with fluorescent probes (Gorelick *et al*., 2019). FIB-milled samples (lamellae) do not display compression, ridges and crevasses, and there are no apparent discontinuities in large-scale structural elements inside cells and tissue. However, samples may still exhibit uneven thickness (curtaining, (Rigort *et al*., 2012)), and recent studies have shown that the ion beam used for milling damages the surfaces layers of the milled lamellae up to a depth of 60 nm (Berger *et al*., 2023; Lucas & Grigorieff, 2023). Furthermore, unlike CEMOVIS, most of the sample is lost during FIB-milling, leaving only the lamellae to be imaged. Finally, the size of a lamella is limited to a few square microns (Villa *et al*., 2013; Rigort *et al*., 2012), compared to tens to hundreds of thousands of square microns of a ribbon of CEMOVIS sections. The larger imageable area of CEMOVIS samples is particularly valuable in the study of tissue, which includes sections with multiple cells to address questions that go beyond the confines of a single cell. Therefore, while there are clear advantages to FIB-milling, it also has a number of fundamental limitations compared to CEMOVIS.

In the present study, we sought to assess the damage on the molecular level in CEMOVIS samples. Using 2-dimensional template matching (2DTM), it is possible to measure the degree of integrity of detected targets in the sample, such as ribosomal subunits (Lucas & Grigorieff, 2023). We prepared CEMOVIS section ribbons of high-pressure frozen *Saccharomyces cerevisiae* cells and measured the signal-to-noise ratio of detected 60S ribosomal subunits. Our results demonstrate that 60S subunits remain structurally well preserved in most parts of the sample, although some areas show signs of more extensive damage.

## 2. Results

We prepared vitrified samples from high-pressure frozen *S. cerevisiae* cell paste, cut into sections of nominally 100 nm thickness (Figs. 1(a) and 1(b)). Initial attempts to image these samples showed clear movement of the sections under the electron beam, presumably due to incomplete attachment of some of the sections to the grid surface. To reduce this beam-induced motion, we coated the grids with a 10-nm layer of platinum before sections were transferred to the grid, thereby increasing the percentage of images with no noticeable motion. At higher magnification (calibrated 1.17 Å/pixel), ridges and crevasses in these sections are clearly visible as bands of dark and light areas, respectively (Figs. 1(c) and 1(d)). Some images also show cell and organelle membranes with discontinuities, similar to previous observations (Fig. 1(d)). We collected 933 micrographs and processed them using *cis*TEM (Grant *et al*., 2018).

**Figure 1.**
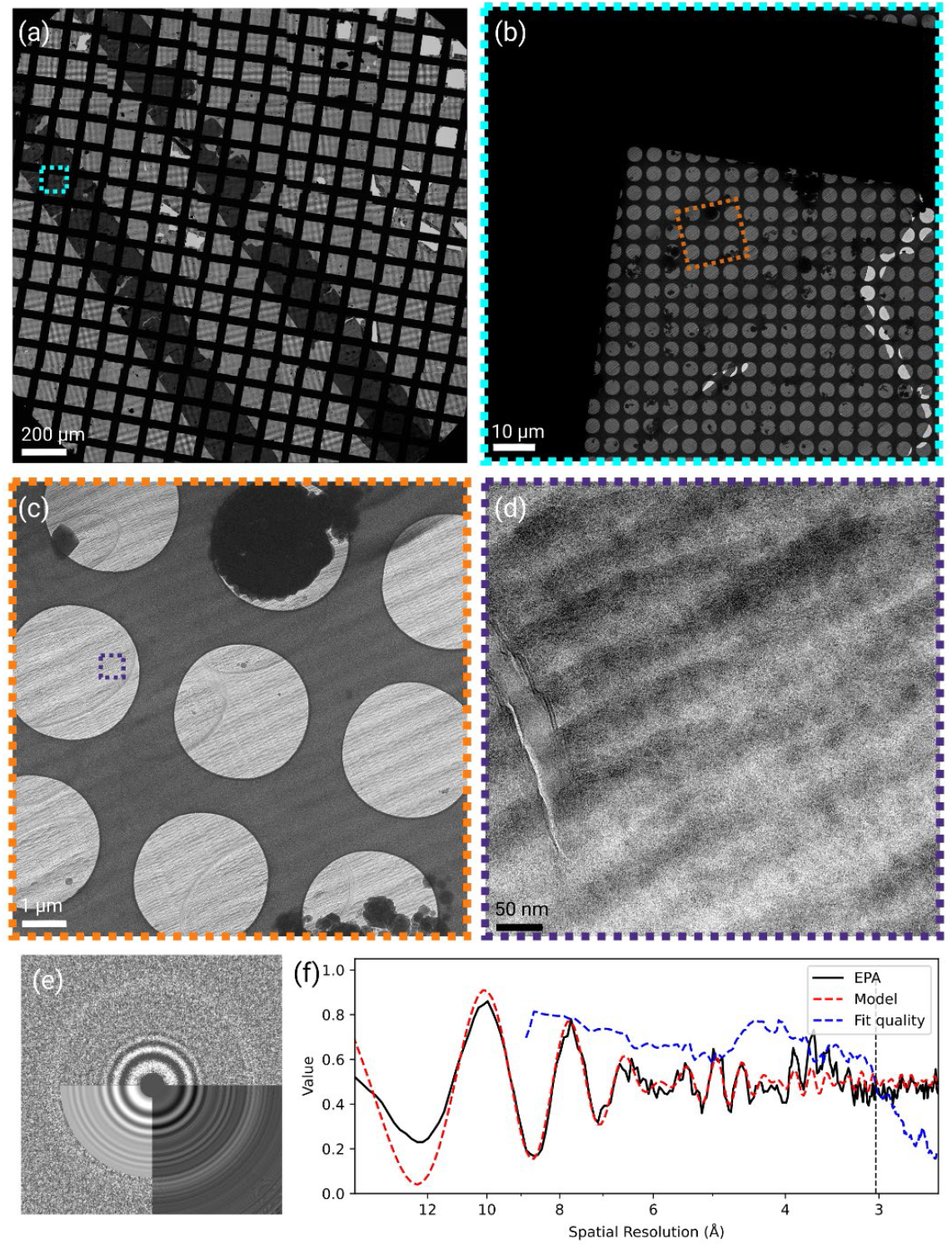
**(a)** Low-magnification image of a 100-nm ribbon of CEMOVIS sections. **(b)** Medium-low magnification view of a grid square covered by a CEMOVIS ribbon. **(c)** Medium-magnification view of carbon film holes covered by a CEMOVIS ribbon. **(d)** High-magnification micrograph of yeast cells within a CEMOVIS ribbon. **(e)** 2D power spectrum of a micrograph shown in (d), overlaid with the equiphasic average (lower right quadrant) and fitted CTF model (lower left quadrant). **(f)** Plot of the equiphasic average of the power spectrum of the micrograph shown in (d), together with the fitted CTF model and “goodness-of-fit” indicator. The fitted parameters show an average defocus of 760 nm and a sample thickness of 174 nm. Thon rings could be fitted up to a resolution of 3.1 Å.

We selected 307 micrographs that showed little or no beam-induced motion based on the trajectories determined during motion correction (Grant & Grigorieff, 2015), and an estimated sample thickness of 150-200 nm based on the Thon ring patterns calculated from the frame averages (Elferich *et al*., 2024) (Figs. 1(e) and 1(f)). Using an atomic model of the 60S subunit of the *S. cerevisiae* ribosome (PDB: 6Q8Y) to generate a template, we searched these images for 60S subunits using 2DTM (Lucas *et al*., 2022). To assess template bias in subsequent 3D reconstructions calculated from the detected targets, we removed atoms from the atomic model within a cubic volume with a side length of 40 Å in the center of the 60S subunit as well as atoms belonging to ribosomal protein L34A (Figs. 2(a) and 2(b)) (omit template, (Lucas *et al*., 2023)). Our search yielded 28,238 60S targets above the standard significance threshold, which is set to allow an average of one false positive per micrograph (Figs. 2(c), 2(d), 2(e) and 2(f), (Rickgauer *et al*., 2017)). Fig. 2 summarizes the search results, comparing number of detected targets (Figs. 2(g) and 2(h)) and the observed 2DTM z-score (Fig. 2(i)) or 2DTM SNR (Fig. 2(j)) values with previous results obtained from FIB-milled lamellae (Lucas *et al*., 2022). The comparison shows that the median 2DTM z-score and SNR values in CEMOVIS sections are lower compared to lamellae (Figs. 2(i) and 2(j)). This may be due to residual beam-induced motion in the CEMOVIS sections, which cannot be completely excluded as a factor in our experiments, as well as the 10-nm platinum coating. The lower number of detected targets in CEMOVIS sections is discussed below. Despite the lower SNR and detection numbers, the 3D reconstruction (Figs. 3(a) and 3(b)) shows clear high-resolution detail in the region omitted in the template, validating the detection of true targets, and demonstrating that CEMOVIS sections preserve molecular structure at near-atomic resolution. Fourier-shell correlation (FSC) plots calculated within the central cube omitted in the template suggests a resolution between 3.1 – 3.3 Å (Fig. 3(c)). When inspecting the density, we found that in the central cube density for RNA bases were well separated, indicative for a resolution better than 3.5 Å. Density for the omitted L34A subunit, located at the periphery of the 60S subunit, was also well resolved (Figs. 3(d) and 3(e)) and we estimated the density to be at 3.5 Å resolution, since larger sidechains were resolved (Fig. 3(f)).

**Figure 2.**
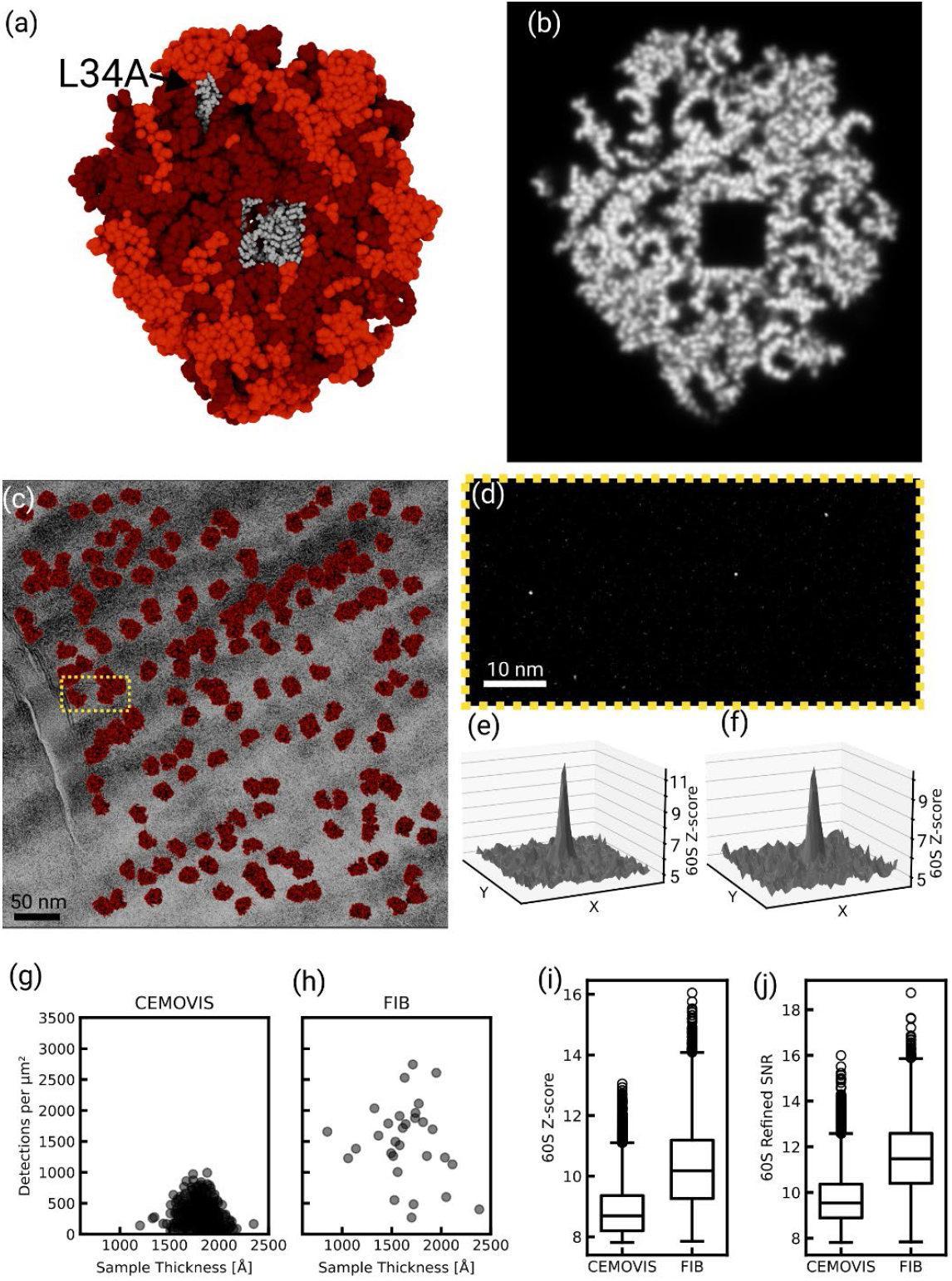
**(a)** Rendering of the 60S model used as a template for 2DTM. Omitted atoms are shown in grey. **(b)** Slice through the simulated density. **(c)** The micrograph shown in Fig. 1(d) overlaid with 2DTM detections of the 60S subunit. **(d)** Magnified region of the cross-correlation maximum intensity projection (MIP), showing three distinct peaks. **(e**,**f)** 3D plot of regions of the MIP around two of the peaks shown in (d). **(g**,**h)** Number of 60S detections per imaged area in micrographs from CEMOVIS (g) and FIB-milling (h) sections plotted against the sample thickness estimated from CTF fitting. **(i)** Box-plot of z-scores after the initial search with the 60S template in CEMOVIS or FIB-milled samples. **(j)** Box-plot of SNRs after refinement of 60S detections in CEMOVIS or FIB-milled samples.

**Figure 3.**
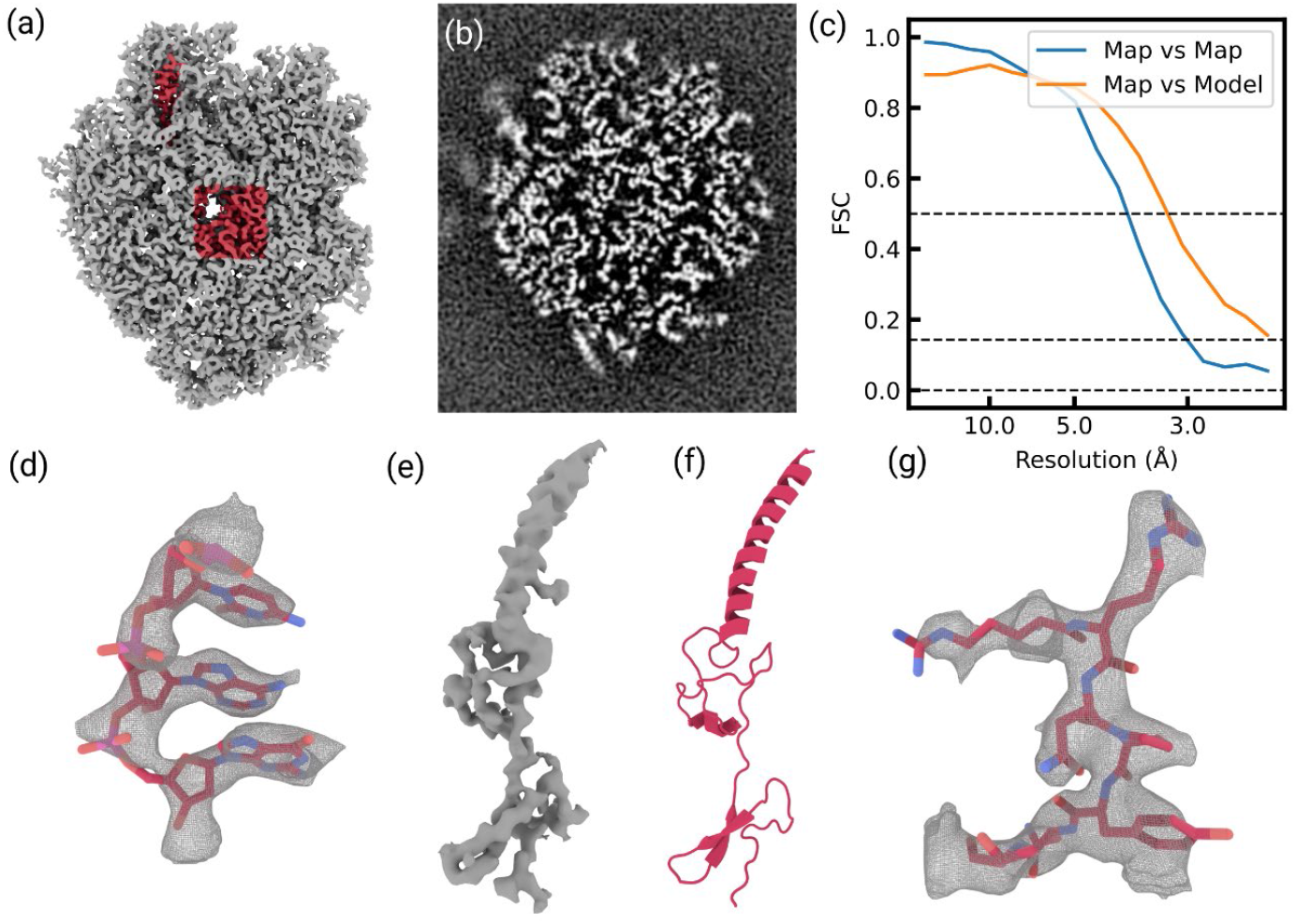
**(a)** Isosurface rendering of a 3D reconstruction from 60S detections in CEMOVIS sections. Parts of the density that were included in the template are shown in grey; parts that were omitted are shown in red. **(b)** Slice through the density of a 3D reconstruction from detections of 60S subunit in CEMOVIS. Compared with Fig. 2(b), there is density in the areas that were omitted from the template. **(c)** FSC calculate for the density in the central box of the map that was omitted from the template. The FSC between half-maps drops below 0.143 at 3.1 Å, while the FSC between map and model drops below 0.5 at 3.3 Å. **(d)** Close-up render of residues 2812 – 2814 of chain BB (25S ribosomal RNA) in PDB ID 6Q8Y, together with an isosurface of the reconstructed map, filtered to 3.2 Å and sharpened with a B-factor of -75 Å^2^. The isosurface mesh was masked at a distance of 2 Å from the shown model. **(e)** Isosurface render of the density attributed to subunit L34A. The map was low-pass filtered at 3.5 Å and sharpened with a B-factor of -30 Å^2^. The mesh was masked at a distance of 2 Å from the model of L34A. **(f)** Cartoon representation of subunit L34A. **(g)** Close-up render of residues 9 – 15 of chain BN (ribosomal protein L34A) in PDB ID 6Q8Y, together with an isosurface of the reconstructed map, filtered to 3.5 Å and sharpened with a B-factor of -75 Å^2^. The isosurface mesh was masked at a distance of 2 Å from the shown model.

We also attempted to determine the dependence of 2DTM SNR on the depth inside the sample (z-coordinate). In FIB-milled samples, there is a clear attenuation of SNR values near the sample surface due to FIB-milling damage (Lucas & Grigorieff, 2023) (Fig. 4(a)). However, we did not observe a clear profile in CEMOVIS sections, where 60S detections with low SNR values were apparent within the sample slab (Fig. 4(b)). We then questioned whether the dark and light bands visible in the micrographs, which we assumed to be crevasses, might be correlated with damage. To investigate this, we band-pass filtered micrographs to accentuate the appearance of these bands and plotted the location of 60S detections (Figs. 4(c) and 4(f)). 60S detection with low or high SNR scores were visually apparent in both dark and light areas of the micrograph and this observation was supported by the similarity of the distributions of the overall pixel intensity variation and pixel intensity at 60S detections (Figs. 4(e) and 4(h)). However, the distribution also suggested that 60S detections occurred along lines parallel to the dark and white bands, perpendicular to the cut direction. To quantify this behavior, we determined the angle of the crevasses relative to the image x-axis, Ψ_*Crevasses*_, by analyzing the direction of maximal signal variation in band-pass filtered micrographs. We also determined if 60S detections occurred in clusters along the same direction using a modified Ripley’s K function that employs an ellipse instead of a circle. We found in the majority of micrographs a clear angle along which clustering behavior was maximal, which we called Ψ_*Clustering*_ (Figs. 4(d) and 4(g)). Figs. 4(c), 4(d), 4(e), 4(f), 4(g) and 4(h) show in two representative micrographs from two grids with different cutting directions that Ψ_*Crevasses*_ and Ψ_*Clustering*_ coincide. In over 80% of the top 100 micrographs with the highest number of detected 60S, Ψ_*Crevasses*_ and Ψ_*Clustering*_ were identical within 20° (Fig. 4(i)). This suggests that damage to 60S ribosomal subunits is minimal in anisotropic patches that are aligned parallel to the knife edge.

**Figure 4.**
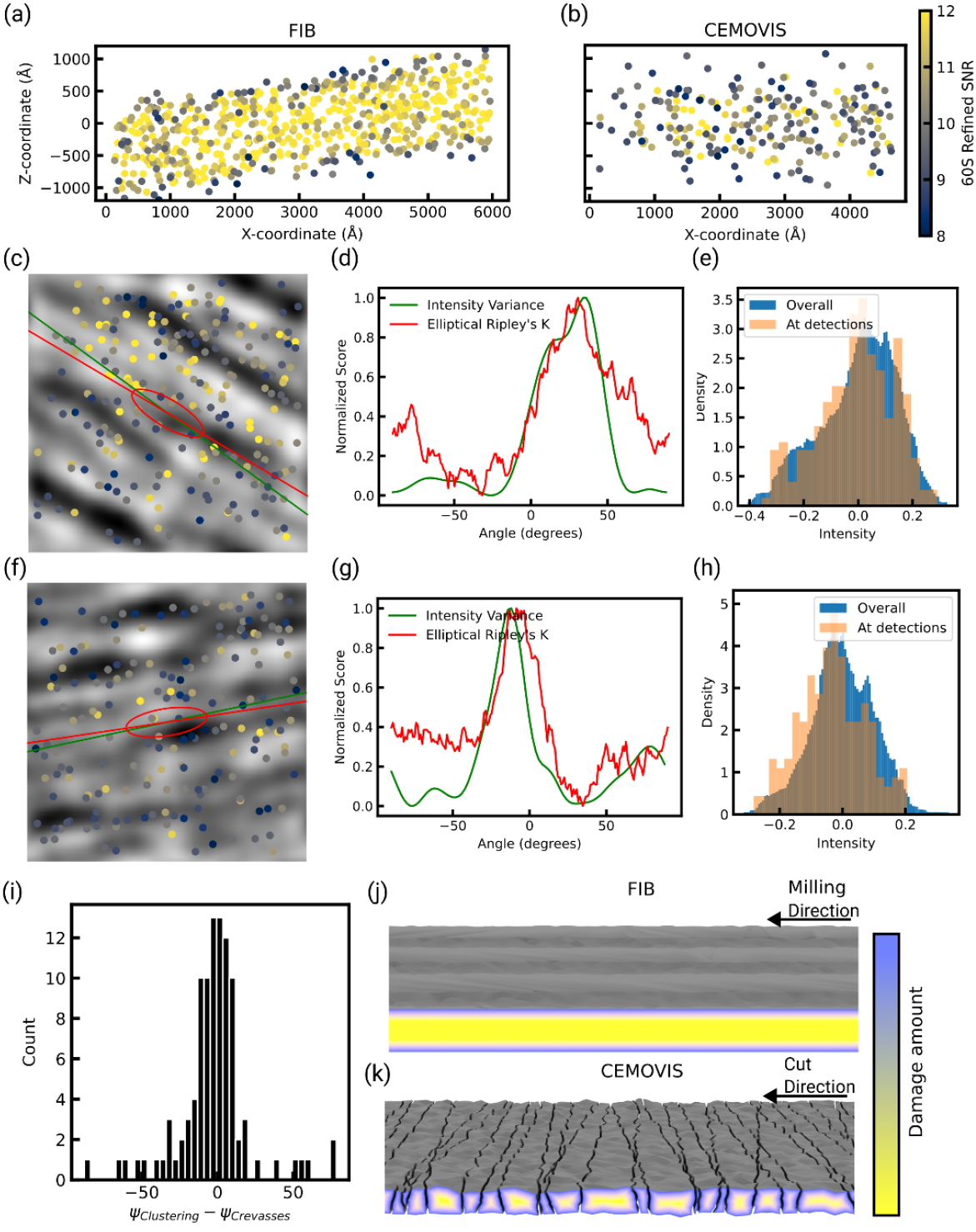
**(a,b)** 60S detections within a representative micrograph from a FIB-milled sample (a) and CEMVOIS sample (a), projected along the y-axis. Points are colored according to the refined SNR scores. Lower scores are found at the top and bottom surface of the FIB-milled sample. No such pattern is apparent in the CEMOVIS sample. **(c**,**f)** Location of 60S detections in two representative micrographs plotted on top of micrographs, which were band-pass filtered to accentuate the crevasses. Points are colored according to the refined SNR scores. A green line indicates Ψ_*Crevasses*_ and a red line indicates Ψ_*Clustering*_. Additionally, the dimensions of the ellipse used for anisotropic Ripleys analysis are indicated as a red ellipse. **(d**,**g)** Plots of the intensity variance score used to determine Ψ_*Clustering* (green)_ and the elliptical Ripley’s K score used to determine Ψ_*Clustering*_. **(e**,**h)** Comparison of the distribution of pixel intensities of the band-passed filtered micrographs shown in (c) and (f) with the pixel intensities at 60S detections. **(i)** Histogram of the difference between Ψ_*Crevasses*_ and Ψ_*Clustering*_ in the 100 CEMOVIS micrographs with the largest number of detections. **(j**,**k)** Schematic of the proposed model for molecular damage in CEMOVIS sections (k) compared to FIB-milled sections (j). Fractures perpendicular to the cut direction cause damage within the slices, constraining detectable 60S subunits to elongated patches parallel to the knife edge.

## 3. Discussion

FIB-milling has become the standard technique to generate thin samples from frozen cells and tissue for cryo-EM and cryo-ET, due to the absence of large-scale sample damage. However, there are a number of downsides to FIB-milling, including the loss of all material removed by the milling process, and molecular damage to a depth of 60 nm from both sides of a lamella (Lucas & Grigorieff, 2023). It is therefore important to investigate alternative techniques for generating thin areas of frozen samples. Here, we revisited CEMOVIS, a technique older than FIB-milling, to assess the molecular damage inflicted by the sectioning process. CEMOVIS samples include many orders of magnitude more area to image, which could lead to a higher throughput of detected targets, and since each section can be imaged it is theoretically possible to image multiple consecutive sections that can then be assembled to a larger 3D volume as previously demonstrated for resin sections (Höög *et al*., 2007). Our analysis of CEMOVIS sections shows evidence of the previously characterized and well-known types of damage, including structural discontinuities and variable thickness resulting from the ridges and crevasses in the sections. However, it remained unclear how the damage affects the integrity of molecules inside the sections. Using 2DTM and 3D reconstruction of detected targets, we obtained direct evidence of structure preservation of 60S ribosomes across larger areas of sections. We did not detect a clear depth profile of structural integrity, such as in FIB-milled lamellae, however, the sections appear to contain bands with more extensive damage that may correspond to cracks generated during cutting (Figs. 4(j) and 4(k)). Nevertheless, a recent study reported an 8.7 Å-resolution protein reconstruction by subtomogram averaging of cryo-ET data of vitreous sections of human brain (Gilbert *et al*., 2024). Additionally, another study demonstrated that vitreous sections of high-pressure frozen lysozyme crystals diffract to 2.9 Å resolution (Moriscot *et al*., 2023). Here we show that 60S ribosomes are preserved in CEMOVIS sections to allow reconstruction at 3.1 –3.5 Å resolution. Interestingly, detected ribosomes tend to cluster along anisotropic patterns parallel to the crevasse direction, but their detection does not correlate with local intensity or thickness variations. This suggests that molecular preservation is influenced more by anisotropic mechanical stresses during sectioning than by overall density or thickness, resulting in elongated patches of higher structural integrity that are not predictable from image intensity alone.

Our experiments highlight some potential improvements of CEMOVIS: To avoid beam-induced motion, it is important to achieve a stable attachment of CEMOVIS sections to the grid surface (Hsieh *et al*., 2006). Currently, sections are attached electrostatically (Pierson *et al*., 2010), which however does not eliminate large variations in attachment. One solution is thus to identify areas of stable attachment by cryo-fluorescence microscopy (Bharat *et al*., 2018). Improved attachment could be achieved by exploring different grid films instead of the commonly used holey or lacey carbon foil. Furthermore, the development of diamond knives with optimized surface modification may reduce cutting artifacts. Finally, thinner sections generally display fewer ridges and crevasses (Al-Amoudi *et al*., 2005), potentially leading to larger areas of uniform structural preservation but also increasing the number of molecules and assemblies partially cut by the knife. Our results indicate that CEMOVIS holds much potential as a technique to image larger volumes of cells, and especially tissue, and that it would benefit from further development to improve samples and reduce preparation artifacts. We propose that the number of detections and average SNR scores after 2DTM using the 60S ribosomal subunit are useful metrics to quantify high-resolution signal preservation when optimizing CEMOVIS protocols.

## 4. Methods

### 4.1. Sample preparation

Haploid *Saccharomyces cerevisiae* of the S288C background (WKY0102; mating type alpha) were grown in YPD at 25°C to mid log phase, pelleted by vacuum filtration (McDonald, 2007) and resuspended in YPD and high-molecular weight dextran to a final dextran concentration of approximately 15% (w/v). The sample was applied to copper tubes and high-pressure frozen using a Leica EMPACT2 (Studer *et al*., 2001). Vitreous sectioning was carried out with a Leica Microsystems EM UC6/FC6 cryo-ultramicrotome equipped with a set of micromanipulators (Studer *et al*., 2014). The instrument was operated at a temperature of -150 °C. A Cryotrim 45° diamond knife (Diatome) was used to trim a pyramid with a height of 40 µm and a side length of approximately 210 µm. Ultrathin sections were produced using a CEMOVIS 35° diamond knife, with a nominal feed of 100 nm, forming a ribbon of sections of 4 to 5 mm in length. The ribbons were placed onto Quantifoil 3.5/1 200 Mesh Cu grids, which had previously been coated with a 10 nm-thick platinum layer using a Safematic CCU/010 HV sputtering device. The thin platinum layer enhances the surface conductivity of the grid and improves the ribbon’s adhesion to the grid. Final attachment of the ribbon to the grid was achieved through electrostatic charging (Pierson *et al*., 2010).

### 4.2. Data collection

Cryo-EM image acquisition was performed using a Krios G4 microscope (Thermo Fisher Scientific) operated at 300 kV in EFTEM mode with a Selectris energy filter and a Falcon 4i camera. The filter slit width was 20 eV. An atlas of the grids was acquired. The selection of sample areas that were well attached to the support film was performed using the Velox software in continuous mode on the Falcon 4i camera at high LM or low SA magnification while the stage was oscillating over a tilt range of ±15°. The subsequent steps were performed using the EPU software. In the Atlas section, a grid square previously identified as containing a well-attached portion of the section ribbon and having a sufficient cell concentration was manually selected. In the Hole Selection section, an image was taken at 470x LM magnification, and a hole in the cell region was chosen. Then, in the Template Definition section, an image was captured at 4800x SA µP magnification, which is suitable for accurate automatic hole detection on the Quantifoil grid used here. On the Captured Image section, about 10 positions for final imaging in the cell regions were selected. The sample focus was manually set to -0.5 µm before starting the data acquisition. High magnification images were acquired at each position with a pixel size of 1.17 Å (105,000x magnification, Nano Probe mode), and an illumination dose of 40 e^−^/Å^2^. The illuminated area was set to 900 nm. Beam shift was used to navigate between each position. This process was repeated for a total of several hundred final images.

### 4.3. Template matching

Movies were motion-corrected using a version of the program *unblur* (Grant & Grigorieff, 2015) that corrects for local motion by alignment of patches (manuscript in preparation). Defocus and sample thickness of micrographs were estimated using *CTFFIND5* (Elferich *et al*., 2024). 60S template density was generated using a model of the 60S subunits from PDB ID 6Q8Y, where the chain BN and all atoms in a cubic volume with a side length of 40 Å around the center of mass were deleted. The program *simulate* (Himes & Grigorieff, 2021) was used to convert the model into a 3D volume with a pixel-size of 1.06 A (FIB) or 1.17 A (CEMOVIS). Template matching was performed using a GPU-accelerated version of the program *match_template* (Lucas *et al*., 2021) using default parameters. Angle, position, and defocus parameters for each detection were refined by maximizing the SNR score (Lucas *et al*., 2021) using a conjugate gradient algorithm.

### 4.4. 3D reconstruction

The 3D reconstruction was calculated using the *cis*TEM program (Grant *et al*., 2018). A single round of manual refinement was run, using the template density used for template matching as a reference, followed by CTF and beam tilt-refinement using the same reference. Fourier-shell correlation curves (FSC) were calculated between volumes generated by cropping the half-maps, and between a model map generated from the 60S subunit without deleting atoms and the central box (35 × 35 × 35 voxels) that was omitted from the template. Isosurface and model renderings were created using the Molecular Nodes plugin in blender (Johnston *et al*., 2025).

### 4.5. Damage Analysis

The angle between apparent crevasses and the image x-axis, Ψ_*Crevasses*_, in each micrograph was calculated by band-pass filtering each micrograph from 2340 to 585 Å using a Butterworth filter. The micrograph was then rotated by an angle *ψ*, pixel intensities of a 2000 × 2000 pixel central box were summed along the x-axis, and the variance of the resulting values was calculated. This was repeated for all values of *ψ* from -90° to 90° in 2° intervals. Ψ_*Crevasses*_ is defined as the angle where the variance was the highest.

The angle of detection clustering, Ψ_*Clustering*_, was calculated using a version of Ripleys’ K function (Dale *et al*., 2002) that uses an ellipse instead of a circle. The elliptical K function was defined as follows:

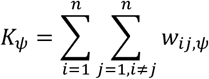

where *n* is the number of detections and *w*_*ij,ψ*_ is 1 if the rotated detection location *R*(*ψ*)r_*j*_ is within an ellipse around *R*(*ψ*)r_*i*_ and otherwise 0. The dimensions of the ellipse were chosen as 600 Å along the x-axis and 200 Å along the y-axis. *K*_*ψ*_ was calculated from -90° to 90° degrees in 1° intervals and Ψ_*Clustering*_ is defined as the angle where the K function reached a maximum.

## Acknowledgements

Data were acquired on an instrument of the Dubochet Center for Imaging in Bern and supported by the Microscopy Imaging Center (MIC) of the University of Bern.

## Data availability

The micrographs collected of the CEMOVIS sections analyzed in the present work were deposited in the EMPIAR database (EMPIAR-XXXXX).

